# Decoding the neural stages from action and object recognition to mentalizing

**DOI:** 10.1101/2025.11.25.690412

**Authors:** Moritz F. Wurm, Seoyoung Lee

**Affiliations:** CIMeC – Center for Mind/Brain Sciences, University of Trento, 36068 Rovereto, Italy; Department of Psychology, University of Chicago, Chicago, IL 60637, USA

## Abstract

Higher-level action interpretation, such as inferring underlying intentions and predicting future actions, requires the integration of conceptual action information (e.g. "opening") with semantic knowledge about persons and objects (e.g. "my friend Anna", "pizza box"). However, how the neural systems for action and object recognition and memory interact with each other to form the basis for inferring higher-level mental states remains unclear. Here we use fMRI-based crossmodal multiple regression representational similarity analysis in human female and male participants to elucidate the processing stages from basic action and object recognition to mentalizing. We show that inferring intentions from observed actions or written sentences involves a modality-general network of lateral and medial frontoparietal and temporal brain regions associated with conceptual action and object representation and mentalizing. The representational profiles in these regions are explained by models capturing different types of conceptual information, revealing distinct but partially overlapping networks for action, object, and mental state representation. There was no strict separation of networks for action, object, and mental state representations, arguing against a sequential bottom-up hierarchy from action and object understanding pathways to the mentalizing network. Rather, left-hemispheric regions, specifically ventrolateral prefrontal, inferior parietal and anterior lateral occipitotemporal cortex, showed strong representational overlap, pointing towards a core network for making meaning of action-object structures at a conceptual level. We argue that this core network represents a distributional semantic hub between classic networks for action and object understanding and the mentalizing network.

**Significance Statement:** How does the human brain integrate information from actions, e.g., "open a pizza box", to understand the actions’ underlying intentions? To do so, the brain needs to combine information from different neural networks—for action and object recognition—and pass them to the mentalizing network for inferring intentions, such as "satisfying hunger". We characterize the interplay of networks using fMRI-based crossmodal multivariate analyses and find that a left-lateralized core network in inferior frontal and parietal cortex and lateral occipitotemporal cortex represents all critical ingredients—conceptual action and object information as well as higher-level mental state representation simultaneously in an overlapping manner. This suggests that this core network is essential for semantic interpretation and functions as bridge between recognition pathways and the mentalizing system.

## Introduction

Understanding others’ actions is central to mentalizing—the ability to infer hidden mental states such as desires, feelings, and intentions. In naturalistic settings, recognizing an action alone (e.g., *opening*) is insufficient; it must be contextualized by integrating semantic information about the people and objects involved. For instance, recognizing someone opening a *pizza box* allows us to infer that she is *hungry* and wants to eat. Importantly, this interpretation builds on the access of action and object knowledge at the conceptual level, regardless of whether the stimulus is visual or verbal. This raises the question how conceptual action and object representations interact with each other to in turn activate associated representations of mental states.

Neural systems for recognizing actions, objects, and for mentalizing have typically been studied separately, and it is implicitly assumed that they are largely anatomically distinct: Action recognition involves the lateral occipitotemporal cortex (LOTC), anterior inferior parietal lobe (aIPL), and premotor cortex (PMC) (Van Overwalle and Baetens, 2009). Left anterior LOTC represents actions at a conceptual, modality-general level (Lingnau and Downing, 2015; Wurm and Caramazza, 2022). Object recognition primarily involves the ventral "what" stream in ventral temporal cortex (VTC) (Goodale and Milner, 1992; Kravitz et al., 2013). While the ventral stream partially overlaps with LOTC, conceptual knowledge about object categories and properties is more prominent in medial temporal cortex (Fairhall and Caramazza, 2013; Clarke and Tyler, 2014; Martin et al., 2018). Mentalizing engages a network comprising the temporoparietal junction (TPJ), dorsal and ventral medial prefrontal cortex (dmPFC, vmPFC), PC, and anterior temporal cortex (ATC) (Frith and Frith, 2006; Van Overwalle, 2009; Thornton and Mitchell, 2018).

Thus, a straightforward model for the pathway from recognizing transitive actions (e.g. opening a pizza box) to mentalizing would be a hierarchy that starts with parallel but separate pathways for action and object recognition. The activated semantic representations associated with the action and object concepts are then integrated to infer possible underlying mental states, possibly drawing on an intermediate stage that bridges the action and object recognition systems and the MN (Fig. 1A). Potential substrates for this integrational stage are lateral mid superior temporal cortex (lmSTC) (Frankland and Greene, 2015), left ventrolateral prefrontal cortex (vlPFC) (Badre and Wagner, 2007; Wurm and Schubotz, 2012), and AG (Binder et al., 2009; Carter and Huettel, 2013; Frankland and Greene, 2020).

**Figure 1.**
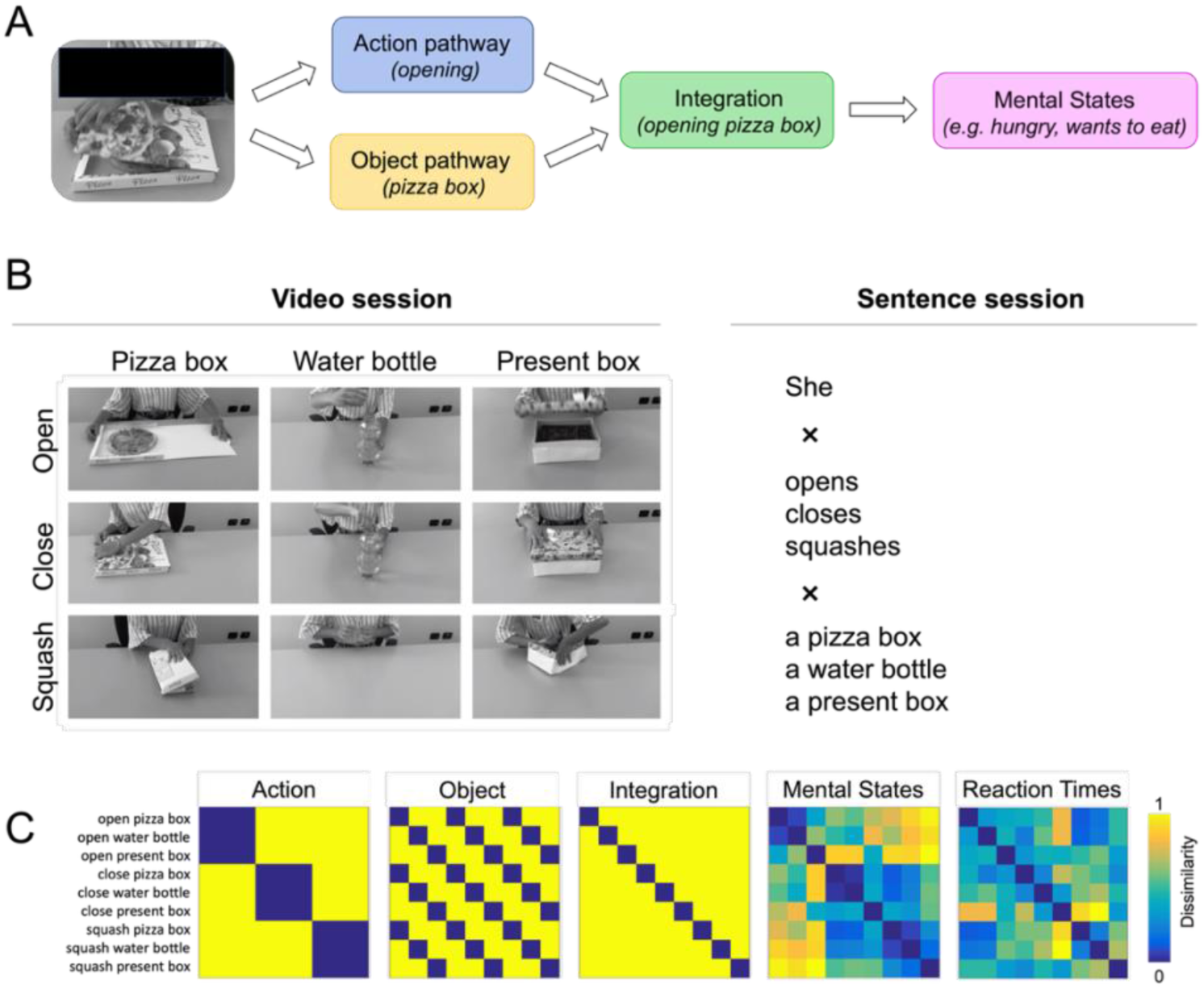
(A) Model for the integration of action and object recognition pathways and the mentalizing network. (B) Experimental design. In two separate fMRI sessions, participants observed videos of actions and corresponding sentences while performing a mentalizing task that requires participants to think about why the action is carried out (i.e., the "why" task) and to press a button as soon as intention is inferred (see Methods for details). (C) Representational dissimilarity models (RDMs) used for multiple regression RSA. The Mental States model was based on ratings targeting intentions, feelings, and broader mental state dimensions associated with the action-object combinations (independent sample of 13 subjects; see Methods for details). The Reaction Time model was based on mean reaction times from the "why" task button presses (mean RTs: 2.62s ± 0.021 SEM for action videos, 2.86s ± 0.017 SEM for action sentences).

An alternative hypothesis is that the systems for action and object recognition and mental state inference are less strictly separated and represent action, object, and mental state information in an overlapping manner. Indeed, conceptual object representations overlap with the MN in angular gyrus (AG)/TPJ, dorsomedial prefrontal cortex (dmPFC), and precuneus (PC) (Fairhall and Caramazza, 2013; Leonardelli and Fairhall, 2022). Conceptual action and object representations overlap in ventral LOTC (Wurm and Caramazza, 2019). The MN and the AON are largely distinct, but there is some evidence for overlap in the right posterior superior temporal sulcus (pSTS) (Deen et al., 2015; Arioli and Canessa, 2019). In addition, the MN and the AON are complementarily activated when people think about *why* as opposed to *how* an action is carried out (Spunt and Lieberman, 2012). To date, no study has investigated the representation of action, object, and mental state information in a unified experiment. It therefore remains unclear whether the overlapping nodes represent heterogenous information or whether the overlap results from co-activation of associated information (e.g. objects might activate associated action concepts in LOTC; mentalizing might draw on semantic object representations in dmPFC, PC, and AG/TPJ).

The present study examined the representational content in the three networks to elucidate the pathway from action and object recognition to higher-level mental state inference. Using fMRI-based crossmodal multiple regression RSA (Wurm and Caramazza, 2019), we localized modality-general representations of actions, objects, mental states, and a putative integrational stage. We found that left inferior frontal and parietal cortex and LOTC represent conceptual action and object information as well as higher-level mental state representation in an overlapping manner, arguing against a strict separation of networks for action and object representation and mentalizing.

## Results

To investigate the processing pathway from basic action and object recognition to conceptual action and object representation and higher-level mental state representations, we used a crossmodal fMRI paradigm in combination with a mentalizing task that required participants to access all these stages of processing. In two separate fMRI sessions, participants observed videos of nine actions and corresponding action sentences (e.g. she is opening a pizza box; Fig 1B) while thinking about why the action is carried out (e.g. because she is hungry and wants to eat). In the video session, responses were correct in 83.3 % of trials (1.4 SEM); in the sentence session, responses were correct in 79.6% of trials (1.6 SEM).

### Modality-general representations during action-related mentalizing

In previous studies, modality-general action representations were found in left anterior LOTC only (Wurm and Caramazza, 2019). The lack of evidence for the involvement of regions in the MN might suggest that representations of associated mental states, such as intentions, were not automatically activated. This may be due to the experimental task, which required action understanding but not higher-level functions such as mentalizing. Hence, we first tested whether action videos and sentences could be decoded beyond the AON, specifically in regions associated with object recognition and memory and mentalizing, when participants engaged in reasoning about the actions’ underlying mental states. We decoded the nine actions both within modality (videos, sentences) and across modality—that is, we trained a classifier to discriminate the activation patterns associated with the observed actions and tested the classifier on its ability to discriminate the activation patterns associated with the corresponding action sentences, and vice versa. Both the decoding of action videos and action sentences as well as decoding across action videos and sentences (Fig. 2, Tab. S2) revealed above chance decoding accuracy not only in brain regions associated with conceptual action representation (left anterior LOTC) (Wurm and Caramazza, 2019), but also in regions associated with conceptual object representation (left posterior middle/inferior temporal gyrus (pMTG/ITG), VTC, lateral PFC, dmPFC, PC) (Fairhall and Caramazza, 2013; Leonardelli and Fairhall, 2022) and mentalizing (dmPFC, TPJ, ATC, PC, vlPFC) (Spunt and Lieberman, 2012). Additional above-chance decoding was observed in bilateral aIPL. The strengths of the crossmodal as well as of the video and sentence decodings were comparable to similar studies using decoding across vision and language (Fairhall and Caramazza, 2013; Wurm and Caramazza, 2019). These results show that not only the left anterior LOTC but a much larger network of brain regions represents modality-general information if the task requires higher-level action interpretation beyond action recognition—that is, mental state inference.

**Figure 2.**
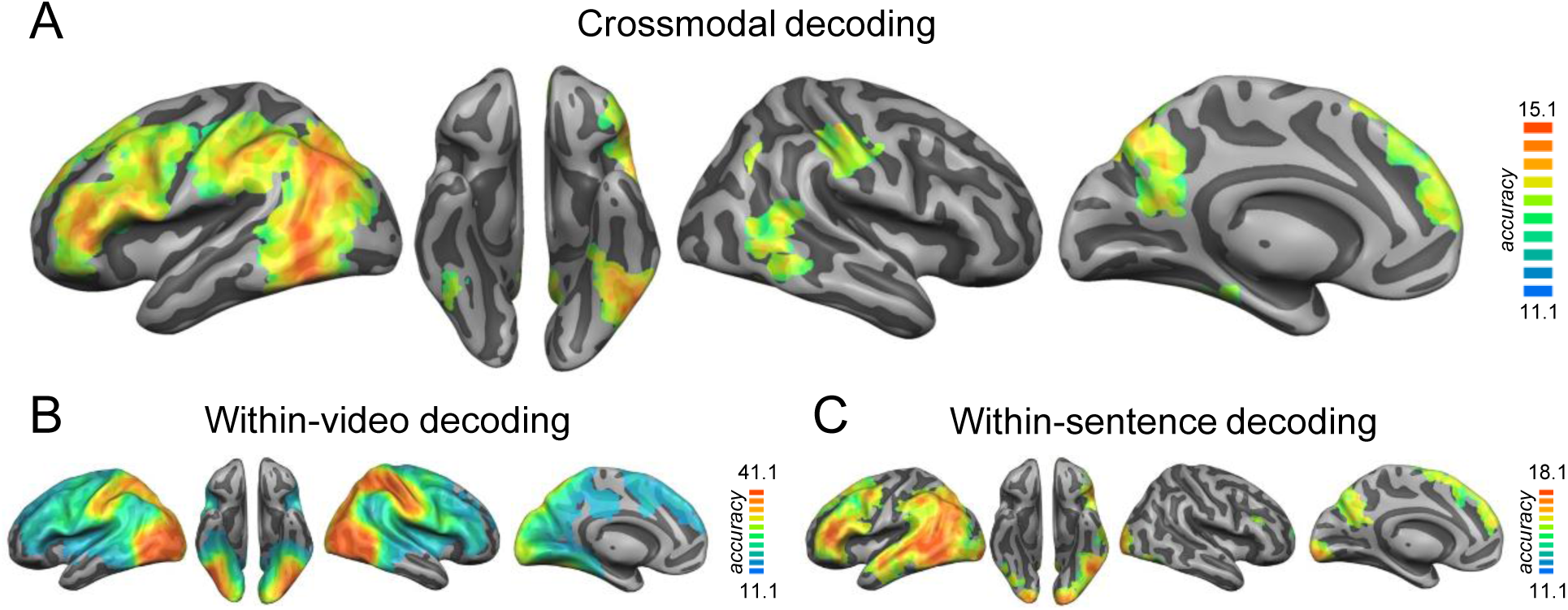
Crossmodal (A), within-video (B), and within-sentence (C) decoding of the nine action-object combinations (decoding accuracy at chance = 11.11%). Mean accuracy maps were thresholded using Monte Carlo correction for multiple comparisons (voxel threshold p=0.001, corrected cluster threshold p=0.05).

### Crossmodal representation of actions, objects, action-object integration, and mental states

Principally, the crossmodal decoding in the obtained regions may be driven by various levels of conceptual representation: actions, objects, the integration of actions and objects, or mental states. To characterize the level of representation with regard to these four stages of processing, we carried out a crossmodal multiple regression RSA: We first extracted from different ROIs the pairwise classifications of the crossmodal decoding described above and converted them into representational dissimilarity matrices (RDMs; see Methods for details). We tested three sets of ROIs associated with (1) modality-general action representation (Wurm and Caramazza, 2019), (2) modality-general object representation (Fairhall and Caramazza, 2013), and (3) mentalizing during action observation / sentence reading (Spunt and Lieberman, 2012). These ROIs also covered candidate regions for the hypothesized integrational stage (left AG, left vlPFC, and left ATC2; see Fig. 3, top right panel). The RDMs extracted from the ROIs were entered into a multiple regression with four models of interest (Fig. 1C): Models for action and object information, an integration model, and a mental states model that captures the similarities of the actions’ underlying mental states (Fig. 1C). The integration model is formally defined as the interaction of the action and the object model: The actions and objects can be conceived as compound stimuli, and the integration model captures the unique combination of each action and object. The mental states model was based on ratings that targeted intentions and feelings that could be associated with the nine actions as well as more general mental state dimensions (Tamir et al., 2016; see Methods for details). In addition, a reaction times model was included to regress out more general effects of cognitive load associated with the mentalizing task.

**Figure 3.**
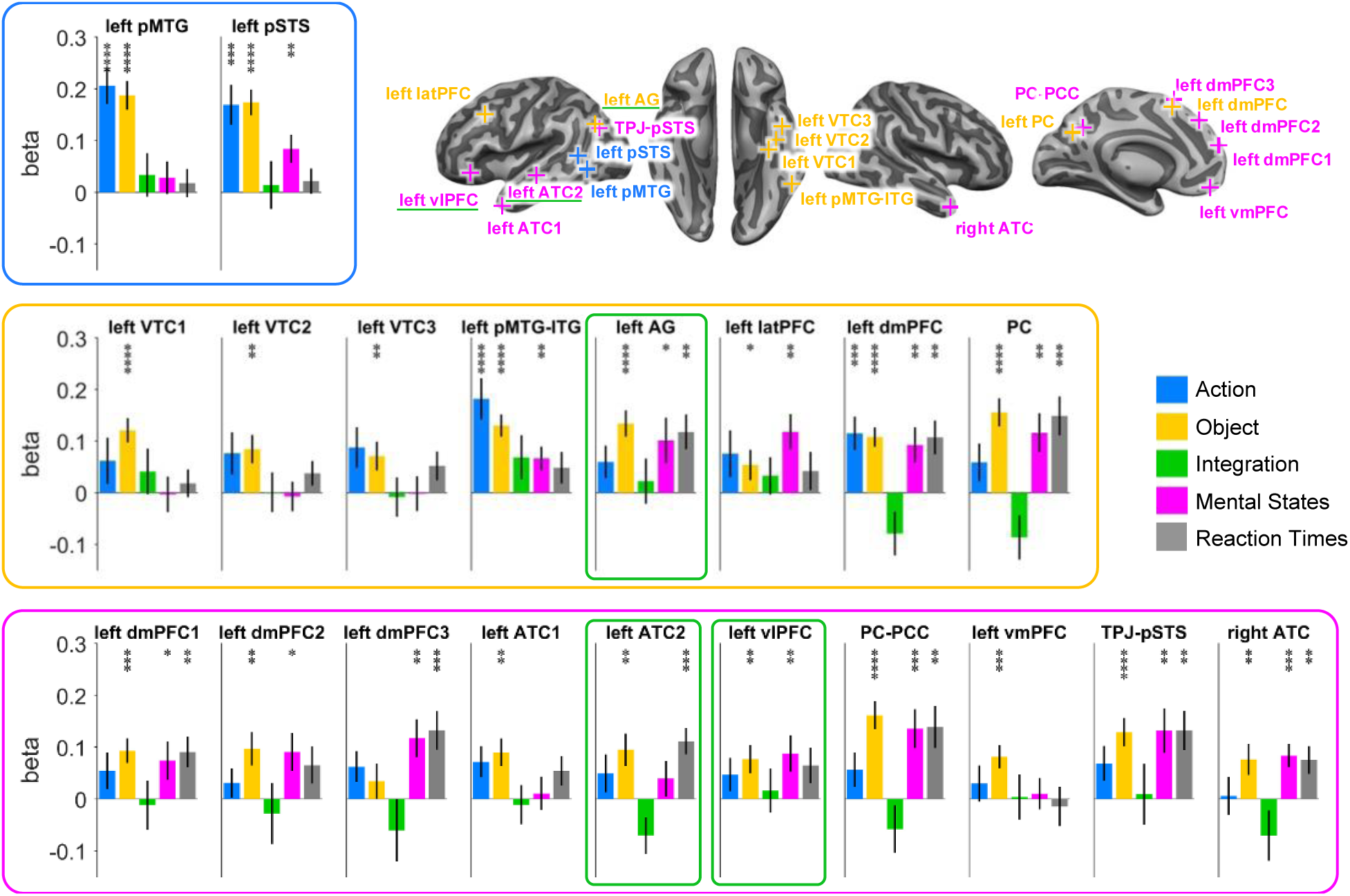
Crossmodal multiple regression RSA ROI analysis. ROIs were based on peak coordinates (upper right panel) for conceptual representation of actions (blue), objects (orange), and mental states (pink; see Methods for details). Hypothesized regions for integrational representations are marked in green. All action ROIs reveal modality-general action-specific representations, all object ROIs reveal modality-general object-specific representations, and seven of ten mental states ROIs reveal modality-general mental state representations. No ROI showed evidence for an integrational stage of processing. Asterisks indicate FDR-corrected significant decoding accuracies above chance (* p<0.05, ** p<0.01, *** p<0.001, **** p<0.0001). Error bars indicate SEM.

We found that each of the three sets of ROIs exhibited the corresponding type of representation: In all action ROIs, the representational content could be explained by the action model; in all object ROIs, the representational content could be explained by the object model; and in seven of the ten mentalizing ROIs (except left and right ATC and vmPFC), the representational content could be explained by the mentalizing model (Fig. 3). The integration model could not explain the representational content in any of the ROIs. The Reaction Times model explained variance in left AG, left dmPFC, PC, and left and right ATC—all regions belonging to the MN. Since the reaction times reflect the effort of inferring the actions’ underlying intentions, a possible interpretation is that the model captures the cognitive load related to intention inference in MN regions. Notably, the effort involved in intention inference is likely representationally distinct from the inferred intentions themselves. Thus, while the Mental State model captures the similarity of the actions’ associated mental states, the Reaction Times model captures how similar the actions are in terms of the difficulty to infer the mental states, which might particularly draw on classic MN regions (e.g. taking the perspective of the acting person).

A repeated-measures ANOVA indicated that the representational content differed between ROIs (Interaction ROI x MODEL: F(76,1672)=1.55, p=0.002). Interestingly, most ROIs were representationally heterogenous, that is, the representational variance in the ROIs could be explained by two or three of the tested models. This representational heterogeneity can be described by the following pattern: The action ROIs—left pMTG and pSTS—were not only sensitive to conceptual action information but also to object information in the videos and sentences. Similarly, all ROIs sensitive to mental states (except dmPFC3) were also sensitive to object information. Thus, almost all ROIs sensitive to objects were also sensitive to either actions or mental states, or both. Only VTC and vmPFC showed significant effects for the object model only. In left pSTS, left pMTG/ITG, and left dmPFC, all three types of representation—actions, objects, and mental states—were found.

To test whether there were additional regions sensitive to modality-general action, object, integration, and mental state representations, we conducted a searchlight analysis (Fig. 4). This analysis revealed results that were consistent with the ROI analysis. First, modality-general action representations were found in left pMTG/pSTS as well as in bilateral aIPL––another key region of the AON––and in left IFG. Second, modality-general object representations were found in all regions previously reported for conceptual object representation (Fairhall and Caramazza, 2013; Leonardelli and Fairhall, 2022), as well as in the left anterior vlPFC and bilateral aIPL. Third, modality-general mental state representations were found in left vlPFC, left premotor cortex, posterior TPJ, and pSTS extending into middle/inferior temporal cortex. The integration model did not reveal clusters surviving multiple comparison correction. The Reaction Times model revealed clusters in left AG and PC, overlapping with the object model. Again, these results demonstrate partially overlapping representations: First, the strongest overlap––between actions, objects, and mental states—was observed in left LOTC. Second, actions overlapped with objects (but not with mental states) in bilateral aIPL. Third, action clusters overlapped with mental states (but not with objects) in left vlPFC.

**Figure 4.**
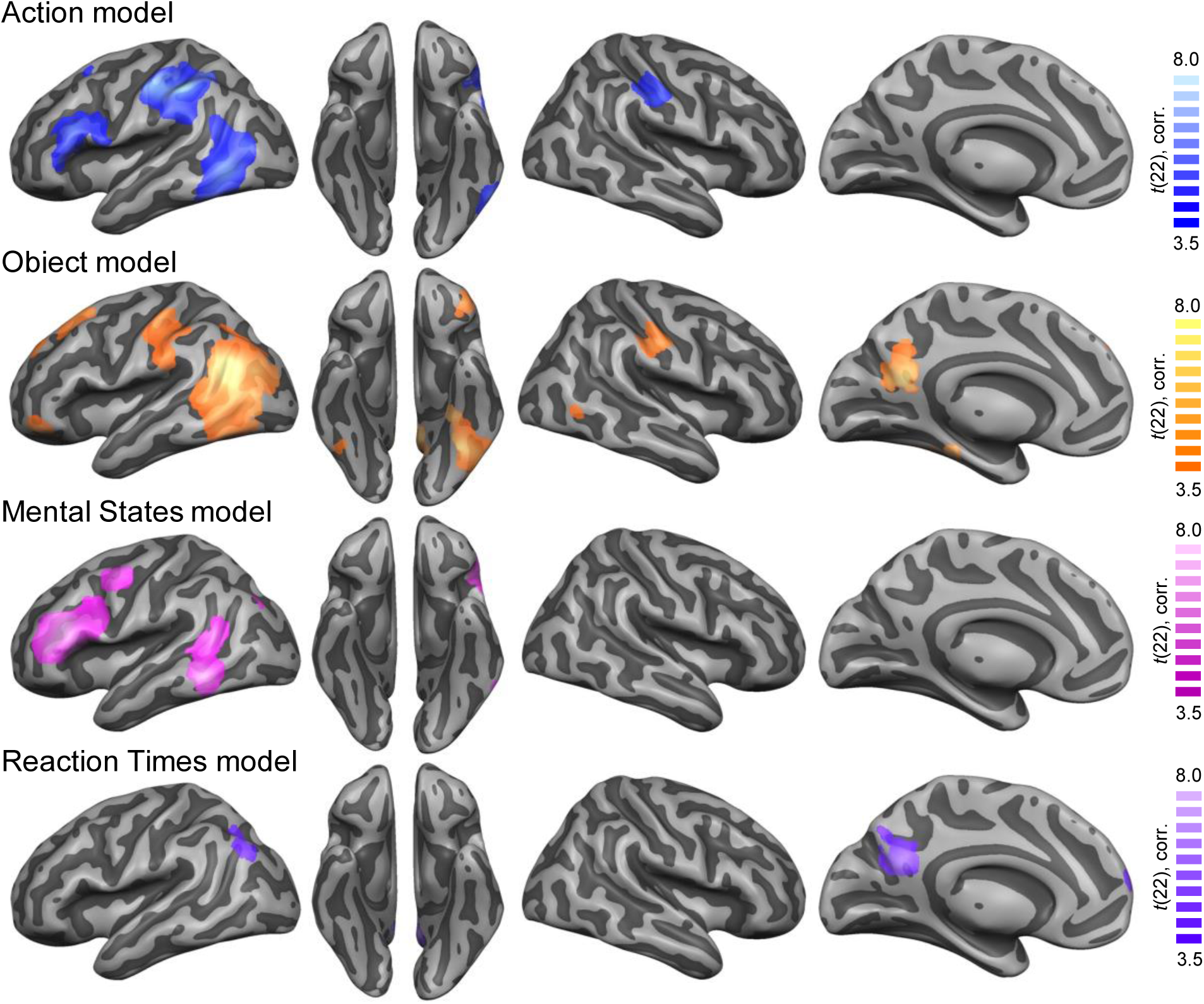
Crossmodal multiple regression searchlight RSA. Maps for action, object, and mental state models revealed overlapping effects in left anterior vlPFC and left anterior MTG and ITG. In addition, overlapping effects for the action and object models were found in left LOTC and bilateral aIPL. No significant effects were found for the integration model. The Reaction Times model revealed clusters in left AG and PC. *T*-maps were thresholded using Monte Carlo correction for multiple comparisons (voxel threshold p=0.001, corrected cluster threshold p=0.05).

Interestingly, the mentalizing cluster in left pSTS/temporal cortex was anterior of the action and object clusters in LOTC, pointing towards a posterior-anterior gradient from action and object representations to a mentalizing-related stage of processing. To characterize this gradient in more detail and show how the representational content changes from posterior occipital to anterior temporal cortex, we performed a path-of-ROI analysis. This was done by plotting the beta values of the five models as a function of position along a vector from posterior middle occipital to anterior middle temporal cortex (Fig. 5). This analysis revealed a significant change from representational content (ANOVA interaction ROI x MODEL: F(156,3432) = 4.15, p < 0.0001). Specifically, conceptual object representations peaked at the level of posterior LOTC, followed by conceptual action representations peaking in anterior LOTC. The peak of mental state representations was located even more anteriorly in the posterior/mid temporal cortex. Finally, we observed a peak for the Reaction Times model slightly anterior of the mental states peak. In line with a recent proposal for an action recognition pathway in left LOTC (Wurm and Caramazza, 2022), these results demonstrate a partially overlapping progression from object to action to mental state representation from lateral occipital to temporal cortex.

**Figure 5.**
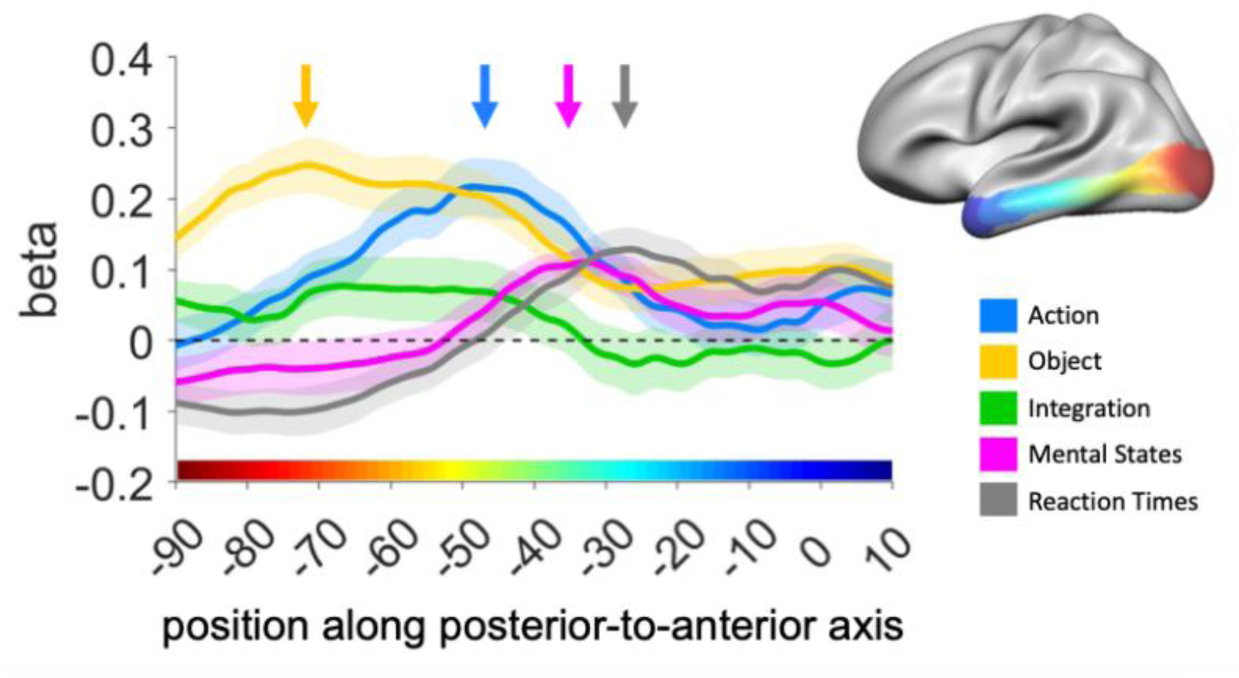
Path-of-ROI analysis. Mean beta values from the crossmodal multiple regression searchlight RSA were plotted as a function of position along the posterior-anterior axis (MNI y-coordinates). The x-axis color bar corresponds to the color coding of the path-of-ROI shown in the brain inset.

### Informational connectivity between action, object, and mental state regions

The multiple regression RSA revealed a complex pattern of partially overlapping modality-general action, object, and mental state representations in the action, object, and mentalizing networks. How do the brain regions of these networks exchange information and thereby transform their representational content? To investigate how the representations in the tested brain regions are related to each other, we conducted an informational connectivity analysis (Coutanche and Thompson-Schill, 2013): We correlated the crossmodal RDMs of the action, object, and mentalizing ROIs with each other, which reflects how distant the representational contents in the different regions are to each other (Fig. 6A). Then, we used multidimensional scaling of the ROI-to-ROI similarities to visualize the representational distances between ROIs (Fig. 6B). In line with the multiple regression RSA, we found that the representations in the action, object, and mentalizing networks partially overlap with each other rather than forming clearly separable networks. This is informative because it allows us to assess where in the brain the networks representationally overlap and where they do not, which might indicate the flow of information between these networks during mental state inference. We observed that all left lateral regions formed a prominent cluster with high representational similarity, distinct from the medial ROIs (PC, dmPFC, vmPFC) and right ATC, which were representationally more diverse. The left-hemispheric cluster groups together nodes from all three networks, including three nodes from the MN: vlPFC, TPJ-pSTS, and ATC. This suggests that information from the action and object networks flows to the MN via these left-hemispheric regions and then onward to other nodes of the MN (i.e., to the midline regions PC, dmPFC, and vmPFC). Due to the high similarity within the left lateral cluster, it is difficult to assess further distinctions, specifically between vlPFC, TPJ-pSTS, and ATC––the three brain regions hypothesized to be likely entrance nodes to the MN. The high similarity between these three MN regions and the action and object nodes (VTC, pMTG, pSTS) suggests that these regions form a common network during action intention inference rather than gradual pathways from one network to the other as illustrated in Fig. 1A.

**Figure 6.**
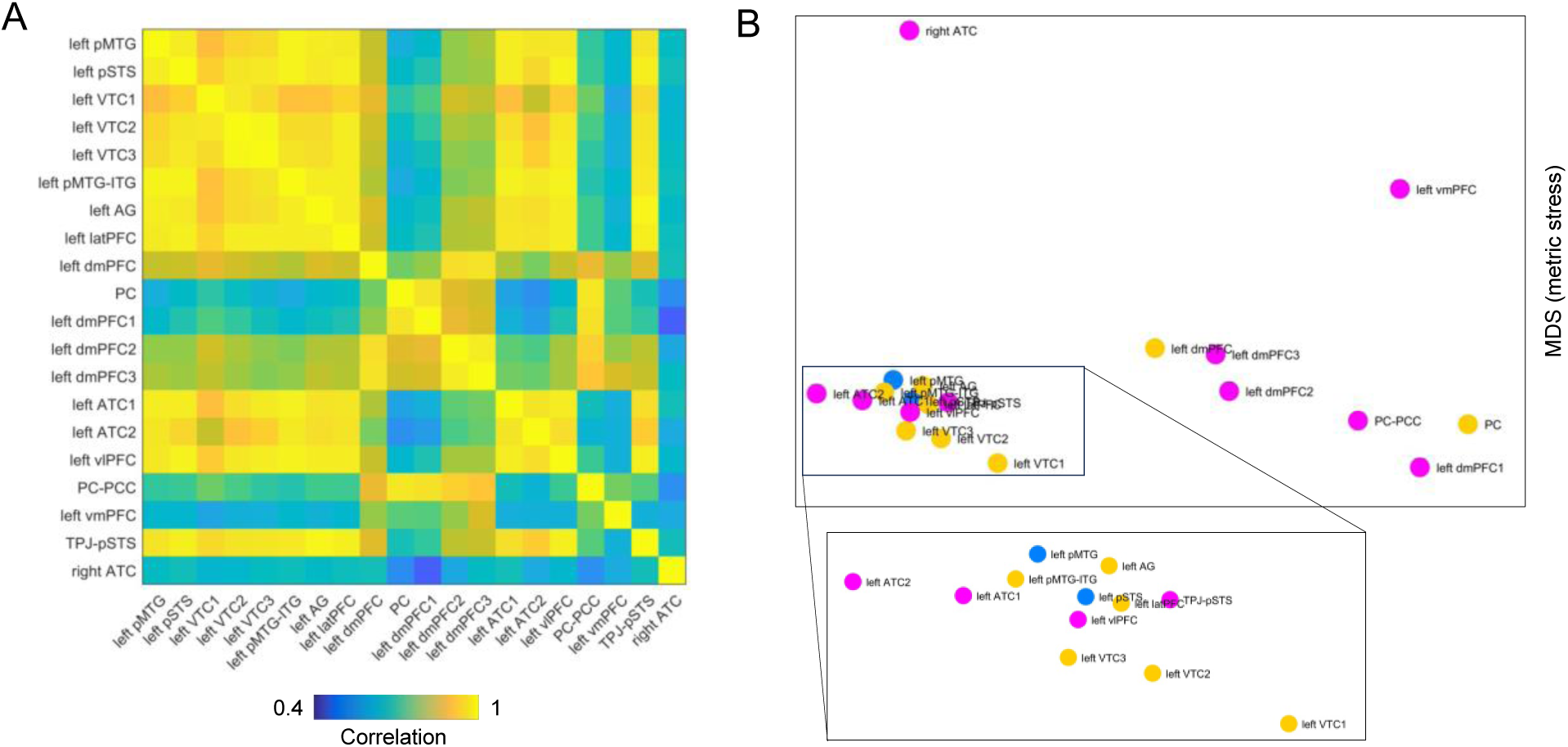
Crossmodal informational connectivity analysis. (A) Correlation matrix from pairwise correlations between group-averaged neural RDMs extracted from the crossmodal decoding. (B) Multidimensional scaling (MDS) of the representational distances between ROIs. Dot colors indicate associated networks for conceptual representation of actions (blue), objects (orange), and mental states (pink).

We also repeated the ROI analysis using ROIs based on cluster peak coordinates from the crossmodal decoding map (Fig. 7A). The crossmodal multiple regression RSA corroborated the results of the literature-based RSA and additionally found left aIPL to represent actions, objects, and mental states (Fig. 7B). Also the multidimensional scaling revealed converging results, demonstrating high similarity between left pMTG, vlFPC, PMC, aIPL, and left VTC as opposed to the representationally more distinct MN regions (left and right TPJ, PC, dmPFC) and right VTC (Fig. 7C and D).

Notably, the high similarity between the three MN nodes (vlPFC, TPJ-pSTS, and ATC) and action and object nodes (pMTG, pSTS, and VTC) is unlikely due to anatomical proximity as the regions are relatively distant. To test this more directly, we computed pairwise Euclidean distances between the ROIs using their MNI coordinates and displayed the distances using multidimensional scaling For both literature-based and decoding-based ROIs, no clustering of the aforementioned regions was observed (Fig. S5). We also correlated the Euclidean distance matrices with the correlation matrices (shown in Fig. 6A and 7C). For literature-based ROIs, the matrices were correlated with each other (r(188)=0.49, p<0.0001), but this effect is likely driven by some highly adjacent ROIs (e.g. VTC1-3, TPJ-pSTS and AG, PC and PC-PCC). Indeed, when we repeated the analysis using decoding-based ROIs (Fig. S5B), no correlation between distance and correlation matrices was observed (r(64)=0.09, p=0.48).

## Discussion

This study aimed to characterize the representational content of brain regions along the pathway from action and object recognition to the MN during mental state inference from actions. We revealed three main findings: (1) When people infer mental states from observed actions or written sentences, conceptual representations are found in a largely left-lateralized network of frontal, parietal, and temporal regions that are typically associated with action and object semantics, mentalizing (Spunt and Lieberman, 2012; Fairhall and Caramazza, 2013; Wurm and Caramazza, 2019), and semantic processing more generally (Binder et al., 2009; Jackson, 2021; Ivanova et al., 2025). (2) The representational profiles in these regions can be explained by models capturing action and object representations (e.g. *open* and *pizza box*, respectively) and mental states (e.g. *satisfy hunger*). We found no evidence for an integrational stage, and there was no strict segregation of action, object, and mental state representations, arguing against a sequential bottom-up hierarchy from action and object recognition pathways to the mentalizing network. Rather, left lateral frontal, parietal, and temporal regions showed strong representational overlap, suggesting that they form a core network for deriving meaning from conceptual action-object structures. (3) Mental state information was found not only in classic MN regions but most prominently in this core network (particularly in left vlPFC, PMC, and anterior LOTC), which likely reflects goal/intention rather than belief inference due to stimulus- and task-related constraints. In LOTC, information is organized along a posterior-anterior gradient from objects to actions to mental states. This might indicate that conceptual action and object information is already transformed to higher-order intention-related information in left LOTC, in interplay with left frontal and parietal regions.

How the neural systems for action recognition and mentalizing relate to and interact with each other has been studied intensively (de Lange et al., 2008; Van Overwalle and Baetens, 2009; Spunt et al., 2010; Becchio et al., 2012; Catmur, 2015). fMRI studies contrasting task demands (e.g. "why" vs. "what" questions) (Spunt et al., 2010; Spunt and Lieberman, 2012) and stimulus conditions (e.g. typically vs. atypically performed actions) (de Lange et al., 2008) typically reveal activation of either the AON or the MN, which can create the impression that these systems are usually complementarily active. However, the hierarchical organization of actions and their underlying intentions presupposes that understanding intentions first builds on action recognition (Kilner et al., 2007; Catmur, 2014), rather than either process being active independently. Our decoding analysis shows that indeed, both the AON and MN are simultaneously active and represent specific modality-general information about the actions if the task requires participants to infer mental states from the actions. By contrast, for tasks that require only action recognition but not mental state inference, decoding of modality-general action representations is largely restricted to left LOTC (Wurm and Caramazza, 2019; Karakose-Akbiyik et al., 2023).

Strikingly, representational content overlaps across networks: The AON regions left anterior LOTC (pSTS and MTG), left PMC, and left IPL also represent mental state information, and the MN regions left TPJ and precuneus represent action information. In addition, almost all MN regions (except right TPJ) represent conceptual object information. For some of these regions, this representational overlap is expected because the three networks anatomically overlap: dmPFC and AG are both nodes of the MN but also of the object semantics network; therefore it might be not surprising that they represent both objects and mental states. The same applies to pMTG/ITG for actions and objects; and pSTS/TPJ/AG for all three types of representation. The present study shows that these regions, while belonging to different networks, not only overlap anatomically but also representationally. It is possible that within these regions, more subtle segregations exist on the individual level. However, this is difficult to assess with multivoxel pattern approaches such as RSA. Representational heterogeneity was observed not only in ROIs where networks anatomically overlap, but also in vlPFC, ATC, aIPL, and anterior LOTC. Among all representationally heterogeneous regions, LOTC/pSTS/TPJ, vlPFC, and aIPL also show high representational similarity (despite their anatomical distance), suggesting they form a core network for integrating action and object information to higher-level mental state interpretation.

While the RSA revealed that the mental states model could successfully capture information in seven out of ten MN regions (and in five out of six decoding-based MN ROIs, except right TPJ), the searchlight RSA showed the strongest effects in regions that differ from the classical MN—namely in left vlPFC/PMC and left anterior LOTC (comprising pSTS, MTG, and ITG), which is associated with semantic reasoning and control (Jackson, 2021; Ivanova et al., 2025). This discrepancy might suggest that inferring mental states from actions does not primarily rely on the classical MN, which is often localized using theory-of-mind tasks, such as understanding false belief vs. physical stories (Saxe and Kanwisher, 2003; Van Overwalle, 2009; Van Overwalle and Baetens, 2009; Spunt and Adolphs, 2014). Indeed, MN regions that showed no effects in the present study, such as right TPJ and vmPFC, are associated with differentiating one’s own beliefs from others’, social decision making, and moral reasoning (Van Overwalle, 2009)—functions that may be less involved in inferring intentions from actions. By contrast, left vlPFC/PMC and anterior LOTC might be involved in retrieving and combining semantic information about actions and objects to infer associated intentions, a function that is usually less required in classic theory-of-mind tasks. Notably, there is often no clear-cut distinction between intentions, understood as abstract, hidden motives for carrying out an action (e.g. *satisfying hunger*), and more concrete, potentially observable goals (e.g. *wanting to eat pizza*) (Csibra and Gergely, 2007; Uithol et al., 2011; Thompson et al., 2019). Accordingly, intention inference may involve the prediction of possible future actions (e.g. that *opening pizza box* is followed by *eating*) (Csibra and Gergely, 2007). It is therefore possible that the representational content captured by the mentalizing model reflects processes important for inferring/predicting future actions. In line with this interpretation, watching videos or reading sentences of predictable vs. unpredictable actions is associated with left vlPFC and MTG (Wurm et al., 2014; Belluzzi and Fairhall, 2025). Thus, we argue that the identified left-lateralized lateral "core network" consisting of anterior LOTC, vlPFC/PMC, and aIPL is critically involved in a more goal-based form of mentalizing that involves semantic reasoning and the inference of possible future actions, as opposed to medial cortex (dmPFC, vmPFC, and PC) and TPJ, which might be more important for belief-based mentalizing that requires theory-of-mind related processes such as belief representation and perspective taking (Schurz et al., 2014; Jamali et al., 2021). This interpretation is supported by the information connectivity analysis, which revealed a representational clustering of left LOTC regions (pSTS, MTG, ITG), left vlPFC, and left aIPL that were representationally distinct from medial MN regions and right TPJ. Goal-based and belief-based mentalizing during action understanding could be dissociated by presenting social scenarios that require the integration of belief information for intention inference (e.g. *giving peanuts to someone knowing vs. not knowing they are allergic)*.

A caveat of this study is the relatively narrow set of nine action-object combinations. To isolate action, object, and mental state information at a conceptual level, we used a conservative crossmodal decoding approach, which generally works better with small numbers of classes, and a controlled design that crossed three actions with three objects. This, however, limited the heterogeneity and complexity of the actions’ underlying mental states. Larger stimulus sets may allow for a better isolation and characterization of goal- vs. belief-based mental state representations. In addition, recent work showed that mental state information in the MN is organized along rationality, valence, and social impact (Tamir et al., 2016)—dimensions that did not substantially vary in the present study. Moreover, inferring an action’s underlying mental states may involve not only the integration of object knowledge but also of knowledge about involved persons, such as age, status, and personality traits (Tamir and Thornton, 2018). For example, we likely infer different intentions from opening a bottle of cleaning detergent if the action is carried out by an adult or a toddler. How person knowledge (and other contextual information) is integrated with action and object information to interpret an action scene in its full complexity is a promising avenue for future research.

In conclusion, the processing from action and object understanding to inferred mental states relies on a left lateralized network of frontal, inferior parietal, and occipitotemporal regions. Each node in this network does not encode unitary information but instead represents multiple types of information simultaneously. This suggests that the nodes largely work in parallel rather than following a strict hierarchical processing from action and object information to inferred mental states. This invites a reconsideration of the strict segregation of recognition pathways (most importantly the AON) and the MN in the context of inferring mental states from actions.

## Acknowledgements

We thank Viola Pegoretti and Leonora Sasso for assistance in preparing the video stimuli and the Caramazza Lab for helpful feedback on the experimental design. This research was supported by the Caritro Foundation, Italy, and the Fondo Italiano per la Scienza.

## Methods

### Participants

27 native Italian speakers (15 females; mean age, 24.2 years; age range, 19-41 years) participated in the fMRI experiment. All participants were right-handed and had normal or corrected-to-normal vision. Four participants were excluded due to bad performance in the task (number of responses during regular trials were less than 2 standard deviations from the mean). The final sample size was therefore N = 23. The study was approved by the Ethics Committee for research involving human participants at the University of Trento, Italy.

### Stimuli

Nine action-object combinations were generated by crossing three actions (open, close, squash) with three objects (pizza box, present, water bottle). For these action-object combinations, video and sentence stimuli were created (Fig. 1B).

For the videos, we filmed each action-object combination performed by two actors (female, male), from two different perspectives (front, side), and using two object exemplars (large, small). In addition, videos were left-right flipped resulting in a total of 16 exemplars per action-object combination (144 action videos in total). Different exemplars for each action-object combination were created to increase the perceptual variance of the stimuli. All 144 videos were edited using iMovie (Apple) and Matlab (Mathworks). The final versions had a length of 2 s, were in gray scale, and a resolution of 400 x 712 pixels.

The sentence stimuli were written descriptions of the action-object combinations. All sentences followed a subject-verb-object structure. To create nine action-object combinations, the verbs chiude, apre, and accartocia (close, open, and squash) were crossed with the objects una bottiglia d’acqua, un cartone della pizza, and una scatola regalo (water bottle, pizza box, and present box). To increase stimulus variance, the verb-object pairs were crossed with six subjects: lei, lui, la ragazza, il ragazzo, la donna, l’uomo (she, he, the girl, the boy, the woman, the man). This created six exemplars per action-object combination (54 sentences in total). Each sentence was presented on a light gray background (400 x 712 pixels) in three consecutive chunks (subject, verb, object), with each chunk shown for 666 ms (2 s per sentence). Different font types (Comic Sans MS, Verdana, MV Boli, Times New Roman, Calibri Light, MS UI Gothic, Arial) and font sizes (32-42) were used to increase the perceptual variance of the sentence stimuli.

Additionally, we created question trials that consisted of questions targeting the possible mental states that could be attributed to the action portrayed in the videos and sentences. These question trials were presented interleaved with the experimental trials (14% of all trials). Each action-object combination was associated with two questions: one designed to elicit an ’agree’ response while the other a ’disagree’ response (e.g. for the video/sentence portraying ’she opens a pizza box’, we expected participants to agree with the question ’does she want to satisfy hunger?’, but disagree with the question ’does she want to be healthy?’). This was to ensure that the participants were paying attention and constantly thinking about the mental states that could be associated with the actions. For each action-object combination and agree/disagree question trial, there were two variants: One focused on intentions (e.g. ’does she want to satisfy hunger?’), whereas the other focused on feelings (’is she hungry?’). Each question was presented on a light gray background (400 x 712 pixels) in one chunk for 2 s using the same font type (Arial) and font size (40).

In the scanner, the stimuli were back-projected onto the center of an acrylic screen via a projector (EPSON EMP-7900) and viewed through a mirror on the head coil (video and sentence presentation 7.5° x 2.4° visual angle). Stimulus presentation, response collection, and synchronization with the scanner were controlled by Matlab with Psychtoolbox extensions (Brainard, 1997).

### Experimental Design

All participants took part in two sessions: a video session and a sentence session. For both sessions, stimuli were presented in a mixed event-related design. Each 2 s video/sentence trial was followed by a 3 s fixation period, which displayed ’perché?’ (why?) on the screen. Participants were instructed to think about the hidden motives of the action, similar to the task by Spunt and Lieberman (2012), which revealed activation of the MN when the ’why’ task was contrasted with the ’how’ task. However, to further motivate the participants to engage in this task, they were asked to press the button on the response box with their right index finger when they had a potential mental state in mind regarding why someone is performing a certain action (e.g. the woman is opening a water bottle because she is thirsty). Participants could respond during the video/sentence or during the ’why?’ fixation period after the video/sentence. During the fixation period, a fixation cross was displayed on the screen for the remainder of the trial, which disappeared as soon as the button was pressed.

Question trials were presented on the screen for 5 s after the presentation of a video/sentence trial. Participants were asked to press one of 4 buttons—from strongly agree to strongly disagree—on the response box. Specifically, they were asked to use the right index finger if they strongly agree with the mental state that was being attributed to the immediate prior action video/sentence, the right middle finger if they partially agree, the right ring finger if they partially disagree, and the right little finger if they strongly disagree. To ensure participants understood the task, a short practice run was completed prior to entering the scanner. To estimate the accuracy of the participants’ responses, we collapsed responses 1 and 2 (generally agree) and responses 3 and 4 (generally disagree).

Trials were presented in blocks of eighteen video/sentence trials, with each action-object combination being shown twice per block and three question trials (21 trials per block in total). Three blocks were presented per run, separated by 10 s fixation periods. Each run began with a 2 s fixation period after five dummy volumes and ended with a 16 s fixation period. Video and sentence runs were presented in separate sessions, with the video session consisting of four functional scans and the sentence session consisting of five functional scans. There were more functional runs for sentences than videos to increase the power for the sentence stimuli, since previous crossmodal decoding studies found that decoding tends to be weaker for sentences compared to videos (Wurm & Caramazza, 2019). The order of the sessions was counterbalanced across participants.

### Data Acquisition

Functional and structural data were collected at the Centre for Mind/Brain Sciences at the University of Trento using a Siemens Prisma 3T MR scanner with a 64-channel phased array head coil. The structural scan was collected between the video and the sentence session. Functional images were acquired using a T2*-weighted gradient echo-planar imaging (EPI) sequence with fat suppression (235 images per functional run; repetition time (TR) of 1500 ms; echo time (TE) of 28 ms; flip angle (FA) of 70°; field of view (FOV) of 200 mm; matrix size of 66 × 66; voxel resolution of 3 × 3 × 3 mm; and 45 transverse slices with a slice thickness of 3 mm in ascending interleaved order). Ten radiofrequency pulses at the beginning of each functional run were sent to avoid T1 saturation. Structural T1-weighted images were obtained using an MPRAGE sequence (176 sagittal slices, TR = 2530 ms, inversion time = 1100 ms, FA = 7°, FOV = 256 × 256 mm, 1 × 1 × 1 mm voxel resolution).

### Preprocessing

Data were preprocessed using BrainVoyager QX 2.8 (BrainInnovation) in combination with the NeuroElf Toolbox, and custom Matlab code. The first volume of the first functional run was aligned to the high-resolution anatomy (six parameters). Data were 3D motion corrected using trilinear interpolation with the first volume of the first run for each participant being used as reference. This was followed by slice time correction. Spatial smoothing was applied with a Gaussian kernel of 8 mm full width at half maximum (FWHM) for univariate analysis and 3 mm FWHM for MVPA.

### Multivariate pattern decoding

Decoding analyses were carried out to identify the brain regions where action-object combinations could be discriminated from each other. For each participant, functional session, and run, a general linear model was computed using design matrices containing predictors for action-object combinations (9 combinations * 2 = 18; two predictors per action-object combination were created by using the first and second half of each run), question trials, the six parameters from 3D motion correction (x, y, z translation and rotation), and 6 temporal drift predictors for high-pass filtering. Each trial was modelled according to the onset and offset of the 2 s video/sentence and these models were then used as reference when fitting the signal time course of each voxel. This procedure created two beta maps per run for each of the nine action-object combinations.

Action-object classification was performed for each participant in volume space using searchlight spheres with a radius of 12 mm and a linear discriminant analysis (LDA) classifier as implemented in the CoSMoMVPA toolbox (Oosterhof et al., 2016). Each voxel was demeaned by subtracting the mean of all betas. Demeaning was executed to remove activations patterns that are common to all action-object combinations.

All classification analyses (crossmodal, within-video, within-sentence) utilized a ’one-against-one’ multiclass decoding approach where each of the nine action-object combinations were discriminated from the remaining eight action-object combinations in a pairwise manner. In the crossmodal action-object classification, a classifier was trained to discriminate the action-object combinations using the data of the video session and its accuracy at discriminating the action-object combinations was tested using the data of the sentence session. This process was repeated conversely (e.g. training the classifier on sentence data and testing on video data) and the resulting accuracies from both classification directions were averaged.

For the within-modality (within-video and within-sentence) action-object classification, a classifier was trained to discriminate the nine action-object combinations using a leave-one-beta-out cross-validation: Seven beta patterns from a total of eight per action-object combination for the within-video classification and nine beta patterns from a total of ten per action-object combination for the within-sentence, were used for training. The accuracy of the classifier at discriminating the action-object combinations was then tested using the held-out data. This procedure was repeated to exhaust all possible training and testing combinations (eight iterations for within-video and ten iterations for within-sentence). The resulting classification accuracies from all possible training and testing combinations were then averaged to produce one classification accuracy per searchlight sphere.

For all decoding schemes, a one-tailed one sample t-test was performed on the accuracy maps to identify voxels in which the decoding accuracy is significantly above chance (11.11%). The resulting t-maps were corrected for multiple comparisons using Monte Carlo correction as implemented in CosmoMVPA (Oosterhof et al., 2016), with an initial threshold of p=0.001 at the voxel level, 10000 Monte Carlo simulations, and a one-tailed corrected cluster threshold of p = 0.05 (z = 1.65). The maps were projected on a cortex-based surface for visualization.

For the multiple regression RSA, the pairwise classifications of the nine action-object combinations were used to create a confusion matrix, which was then converted into a representational dissimilarity matrix (RDM; see below).

### Representational Dissimilarity Models

Action, object, and integration representational dissimilarity models (RDMs) were created by comparing action-object combinations in a pairwise manner and assigning 1 or 0, with 1 representing similarity and 0 representing dissimilarity. For the action model, action-object combinations were similar if they contain the same action irrespective of the object involved. For the object model, action-object combinations were similar if they contained the same object irrespective of the action involved. For the integration model, action-object combinations were similar if they contained both the same action and object (or, in other words, the interaction between action and object model; Fig. 1C).

To generate a mental states model that reflects the similarity of mental states associated with the nine action-object combinations, an independent group of participants (N = 13) took part in a behavioral ratings experiment. In this experiment, sixteen different questions that reflect the potential mental states of the nine action-object combinations were asked for each action-object combination (Supplementary Table 1). The questions were selected to cover a broad spectrum of mental state features: Eleven of the sixteen questions targeted possible intentions and feelings associated with the action-object combinations; the remaining five questions targeted more general mental state dimensions based on Tamir et al. (2016), i.e., valence, arousal, rationality, sociality, and body vs. mind satisfaction. Participants had to respond to every question using a 6- point Likert scale. The mental states model was constructed by computing pairwise correlations between group-averaged responses to the 16 ratings questions for each of the nine action-object combinations. The resulting 9 × 9 matrix was then subtracted from one to create a dissimilarity matrix (Fig. 1C). To test for robustness of the RDM, we computed its split-half reliability. This was done by randomly splitting the sample into two halves (7 and 6 subjects per half), computing the mean RDM for each half, and correlating the RDMs of the halves with each other (using the lower triangle of the matrix, excluding the on-diagonal). This was done in 10,000 iterations; resulting correlation coefficients were averaged across iterations. This resulted in a mean correlation of r = 0.842, which suggests high reliability of the RDM. We also explored more fine-grained models for the three subtypes of mental state features. We created separate models based on the questions on intentions (6 questions), feelings (5 questions), and mental state dimensions (5 questions) using Euclidean distance. These models, which were strongly correlated with each other (r > 0.75) revealed effects that were similar to the grand mental states model (Supplementary Figure S1).

Finally, we included a model for reaction times (RTs) in the multiple regression RSA. This model aimed at controlling for potential differences in cognitive load to infer mental states to the actions. RTs of button presses to the ’why?’ question during action trials were averaged across participants. To construct the model, we first computed the mean RTs for the nine action-object combinations separately for the video and sentence session. The mean RTs were then z-scored for the two sessions to eliminate session-related differences between video and sentence RTs. For the crossmodal RSA, we computed the pairwise Euclidean distances between the nine RTs of the sentence session and the nine RTs of the video session. The model thus reflects how similar the nine action-object combinations are in terms of speed to press a button (broadly reflecting the cognitive load to infer associated mental states) across the two sessions. For the within-session RSA, we computed the pairwise Euclidean distances between the RTs of the nine action-object combinations for each session separately.

The neural RDMs were created by extracting the pairwise classifications (the classification matrix) from each ROI or searchlight sphere of each participant’s crossmodal decoding map, symmetrizing the matrix across the main diagonal, and converting the classifications into dissimilarity measures by subtracting 100 – accuracy. To reduce noise in the neural RDMs, we spatially smoothed them by computing the mean RDM of each voxel and of the surrounding voxels within a radius of three voxels.

RSA usually uses the pairwise similarities of the neural and model RDMs (the off-diagonal values of the RDM), whereas the similarity of each condition with itself (the on-diagonal values of the RDM) is not included. However, for the interaction model, which distinguishes each action-object combination from each other, all pairwise comparisons have the same value (0=dissimilar). Thus, the interaction model would be singular and produce meaningless results in the RSA. A solution to this problem is to include the on-diagonal values of model and neural RDMs in the RSA (Walther et al., 2016). Importantly, because we used cross-validation for the multiple regression RSA, the on-diagonal values have interpretable zero-points and can therefore be included in the RSA without biasing (Walther et al., 2016).

The models were significantly correlated with each other (Supplementary Fig. S2). To test for potential collinearity between the models that might lead to estimation problems in the multiple regression, we computed the condition indices (CIs), variance decomposition proportions (VDPs), and variance inflation factors (VIFs) using the colldiag function in Matlab. CIs assess the sensitivity of parameter estimates to near-linear dependencies among predictors. Values below 10 are generally considered unproblematic. VDPs indicate how much of each coefficient’s variance is associated with a given condition index; collinearity concerns arise when two or more predictors show high VDPs (0.5 or greater) for predictor pairs that show a high CI (> 10). VIFs quantify how much multicollinearity inflates coefficient variances, with values below 5 commonly regarded as acceptable (Belsley et al., 1980). The results of these tests (all CIs < 4, all VDPs < 0.86, all VIFs < 3.6) did not indicate potential estimation problems.

### ROI definition

ROIs were defined in MNI space based on three fMRI studies that aimed at identifying modality-general representations of actions (left pMTG: -55, -62, -2; left pSTS: -50, -25, -23) (Wurm and Caramazza, 2019), objects (left VTC1: -33, -25, -23; left VTC2: -39, -16, -26; left VTC3: -42, -7, -26; left pMTG-ITG: -51, -49, -11; left AG: -48, -70, 31; left latPFC: -48, 20, 40; left dmPFC: -15, 17, 49; PC: -3, -64, 31) (Fairhall and Caramazza, 2013), and action intentions (left dmPFC1: -6, 59, 22; left dmPFC2: -9, 44, 46; left dmPFC3: -6, 23, 64; left ATC1:-48, 11, -35; left ATC2: -57, -22, -8: left vlPFC: -48, 28, -11; PC-PCC: -6, -55, 34; left vmPFC: -6; 50, -17; TPJ-pSTS: -45, -64, 34; right ATC: 51, 8, -29) (Spunt and Lieberman, 2012). It is less straightforward to define ROIs for the integrational stage. However, the aforementioned ROIs also included the hypothesized regions for action-object integration, i.e., left ATC2 (anatomical distance to lmSTC in Frankland & Greene, 2015: 12.8 mm), left vlPFC, and left AG. Therefore, we did not define separate ROIs for the integrational stage. For each ROI, volumetric spheres were created around the peak coordinates, using a radius of 12 mm.

### Multiple regression RSA

To characterize the representational content isolated by the crossmodal decoding, an ROI-based multiple regression RSA using the crossmodal classification accuracies was performed. All pairwise classifications of the action-object combinations were extracted from a ROI to construct neural RDMs for each voxel instead of a single classification accuracy for each voxel. Thus, for each ROI voxel, a 9 × 9 confusion matrix was created, averaged across voxels, made symmetrical across the main diagonal, and transformed into model RDMs by subtracting the classification rate (i.e. the number of predictions for an action-object combination divided by the number of predictions for all action-object combinations) from one. This resulted in one RDM per ROI and participant. The off-diagonals (i.e. the lower triangle of the matrix) and the on-diagonals of the models were vectorized, z-scored, and entered as dependent variables into a multiple regression. The correlation coefficients were Fisher transformed and entered into one-tailed one-sample t-tests, which were False Discovery Rate (FDR) corrected for the number of ROIs included in the regression.

### Searchlight multiple regression RSA

A similar procedure as the ROI-based multiple regression RSA was conducted for the crossmodal searchlight decoding multiple regression RSA. Instead of ROIs, the pairwise classification of the crossmodal action-object decoding was extracted from each voxel in the searchlight sphere, resulting in 9 × 9 confusion matrix for each voxel in the sphere-of-interest and one RDM per searchlight sphere and participant. The RSA procedure described was done for each voxel and the resulting beta values were assigned to the center voxel of each searchlight sphere. The searchlight multiple regression RSA was restricted to voxels that yielded significant above chance accuracies in the decoding analysis, which indicates reliably discriminable information of the action-object combinations (Ritchie et al., 2017). Resulting beta maps were entered into one-tailed one-sample t-tests and corrected for multiple comparisons using Monte Carlo Cluster (MCC) correction as described in the *Multivariate pattern decoding* subsection.

For completeness, we also carried out within-modality (within-video and within-sentence) multiple regression searchlight RSA that were based on the neural RDMs derived from the within-video and within-sentence decoding (see Supplementary Figures S3 and S4). However, because the within-video and within-sentence decoding were not specifically sensitive to modality-general (conceptual) information, they are not well suited to detect the conceptual action, object, integration, and mentalizing stages targeted in this study. The decoding generally exploits the information that most reliably discriminates the nine action-object combinations, which is not necessarily conceptual information. In particular, for the within-video decoding, the nine action-object combinations are discriminable based on various perceptual and kinematic dissimilarities. Thus, the integration model is expected to pick up the specific perceptual and kinematic differences between the nine action-object combinations. By contrast, the crossmodal RDMs reflect the dissimilarities of the action-object combinations at a modality-general level only. Therefore, the crossmodal and within-modality RDMs are likely to reflect different kinds of information, which limits the interpretability of the within-modality multiple regression RSA, in particular for the integration model, and the comparability of the crossmodal and the within-modality multiple regression RSA. Another limitation is that the within-video and within-sentence decoding are each based on 50% of the data, which reduces the power of the within-modality multiple regression RSA.

### Path-of-ROI analysis

To investigate the change of representational content along the posterior-anterior axis from occipital to anterior temporal cortex, we used a path-of-ROI analysis (Konkle and Caramazza, 2013; Wurm et al., 2017; Wurm and Caramazza, 2019). Anchor points were based on the posterior and anterior points of occipital and temporal poles, respectively, which were then connected with a vector on the flattened surface along the middle temporal gyrus. Along this vector, a series of 40 partially overlapping ROIs (12 mm radius, centers spaced 3 mm) was defined. From each ROI, beta values were extracted from the searchlight maps of the crossmodal multiple regression RSA, averaged across voxels, and plotted as a function of position on the posterior-anterior axis.

### Informational connectivity analysis

To test how similar the representational content of the ROIs was to each other, we used informational connectivity analysis (Coutanche and Thompson-Schill, 2013). First, we extracted the neural RDMs of the ROIs from the crossmodal decoding map and correlated them with each other. We only used the lower triangle of the RDMs (off-diagonal pairwise distances between actions) because the on-diagonal correct classifications would confound the informational connectivity analysis with the general strength of decoding in the ROIs (i.e., regions with high crossmodal decoding accuracies show stronger similarity with each other than with regions with lower decoding accuracy). We then converted the resulting correlations between ROIs into distances by subtracting 1 – r, and visualized the pairwise distances between ROIs and decoding schemes using multidimensional scaling (metric stress).

---

**Supplementary Figure S1.**
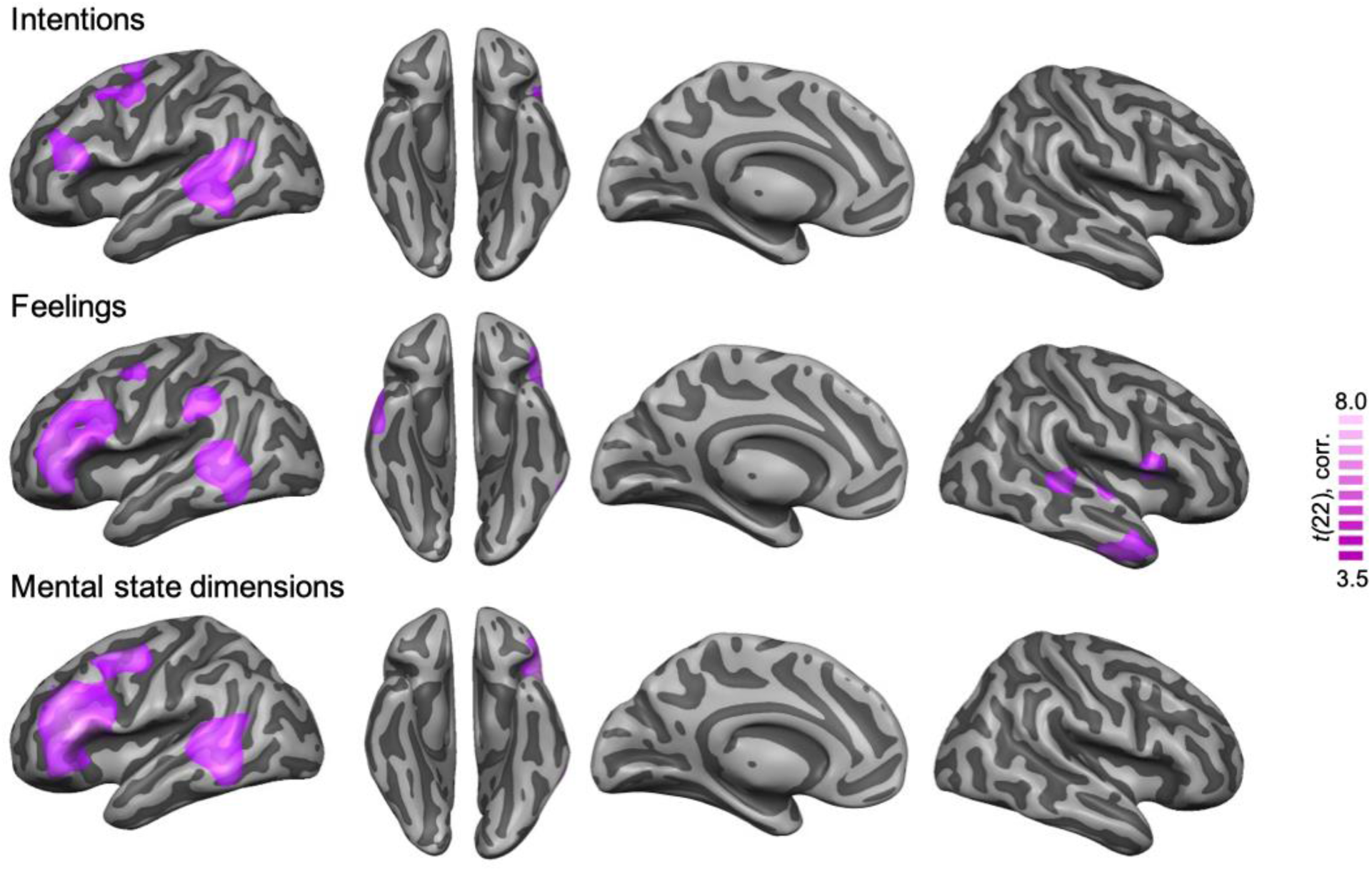
Crossmodal multiple regression RSA for intentions, feelings, and mental state features. The models were based on the questions on intentions (6 questions), feelings (5 questions), and mental state dimensions (5 questions; see Supplementary Table 1). Since the models were strongly correlated with each other (r > 0.75; max. VIF = 11.3, max CI = 11), they were entered in 3 separate multiple regression RSA with the action, object, integration, and RT models. Searchlight maps were thresholded using Monte Carlo correction for multiple comparisons (voxel threshold p=0.001, corrected cluster threshold p=0.05).

**Supplementary Figure S2.**
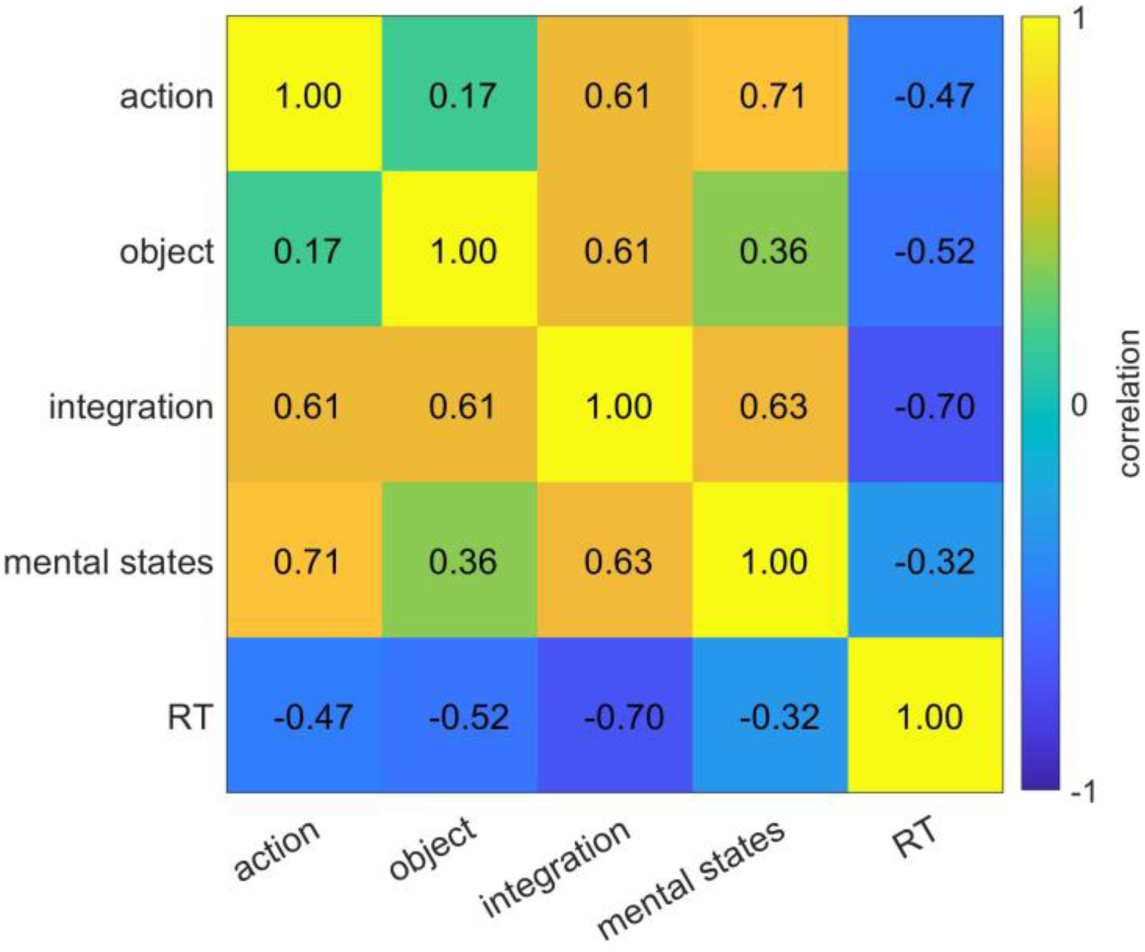
Pearson correlations between the RDMs used for the crossmodal multiple regression RSA.

**Supplementary Figure S3.**
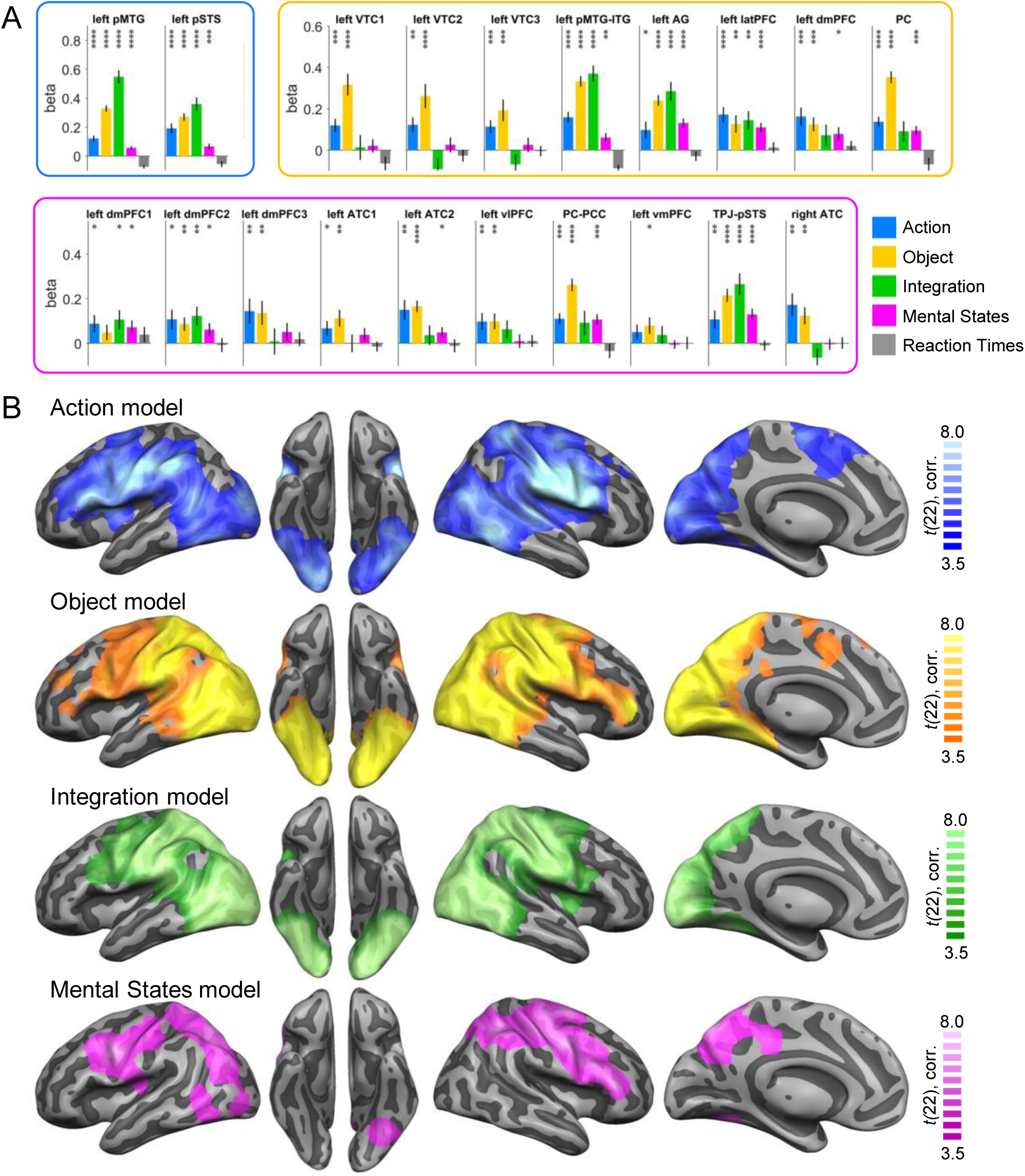
Within-video multiple regression RSA. (A) ROI analysis. Asterisks indicate FDR-corrected significant decoding accuracies above chance (* p<0.05, ** p<0.01, *** p<0.001, **** p<0.0001). Error bars indicate SEM. (B) Searchlight RSA maps. *T*-maps were thresholded using Monte Carlo correction for multiple comparisons (voxel threshold p=0.001, corrected cluster threshold p=0.05). The Reaction Times model did not reveal clusters surviving multiple comparison correction.

**Supplementary Figure S4.**
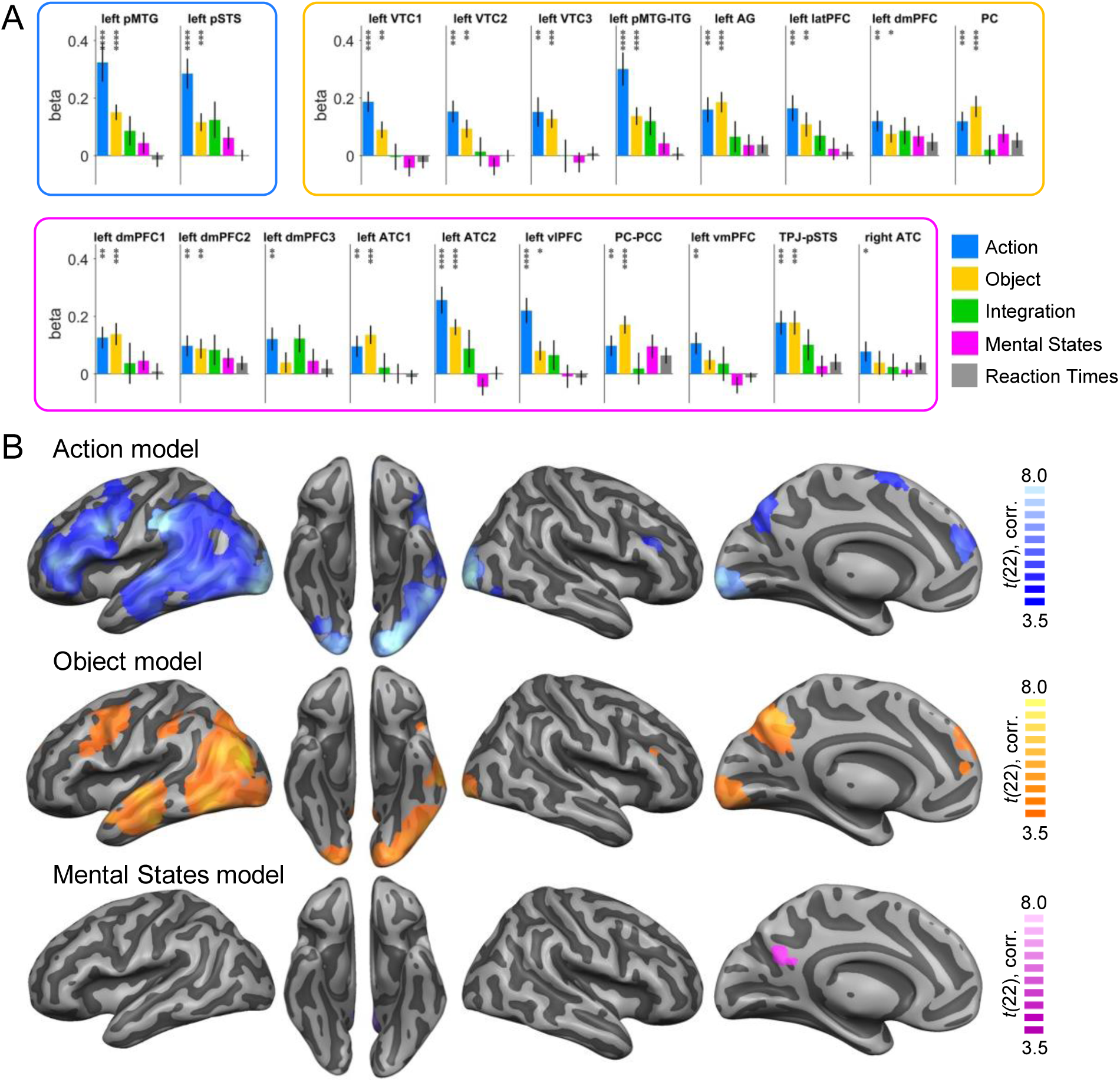
Within-sentence multiple regression RSA. (A) ROI analysis. Asterisks indicate FDR-corrected significant decoding accuracies above chance (* p<0.05, ** p<0.01, *** p<0.001, **** p<0.0001). Error bars indicate SEM. (B) Searchlight RSA maps. *T*-maps were thresholded using Monte Carlo correction for multiple comparisons (voxel threshold p=0.001, corrected cluster threshold p=0.05). The Integration model and the Reaction Times model did not reveal clusters surviving multiple comparison correction.

**Supplementary Figure S5.**
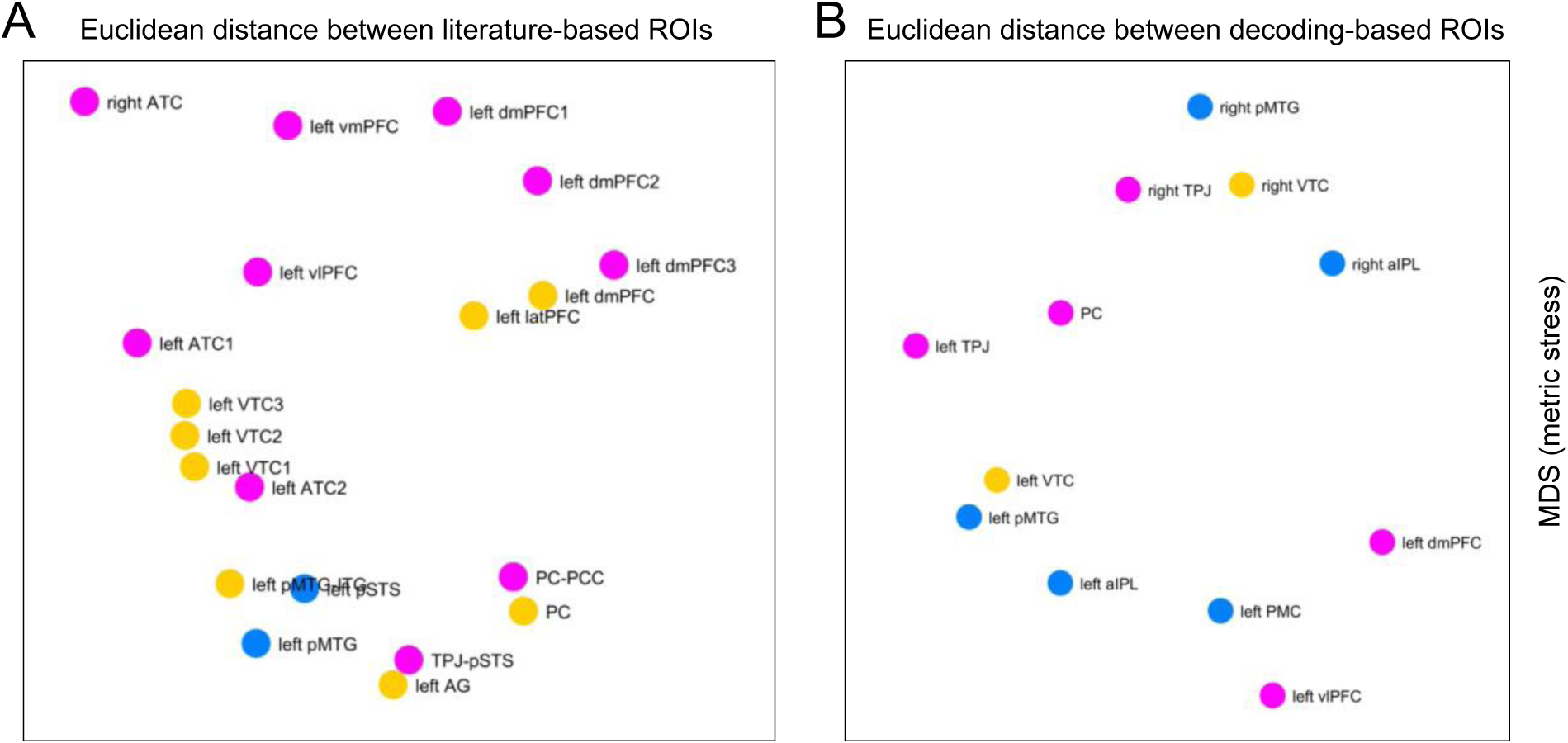
Multidimensional scaling based on Euclidean distance between ROIs. Euclidean distances between MNI coordinates of the literature-based ROIs (A) and decoding-based ROIs (B) were computed, and the resulting distance matrices were entered into multidimensional scaling as reported for the multidimensional scaling based on neural RDMs (see section *Informational connectivity analysis* in the *Methods* section of the main text). Colors indicate associated networks for conceptual representation of actions (blue), objects (orange), and mental states (pink).

**Supplementary Table 1:** Behavioral rating for the mental states model (N=13). Questions 1-6 target intentions, questions 7-11 target feelings of the acting person. Questions 1-11 used a one-sided 6-point Likert scale from 1 (not at all) to 6 (very much) and were selected to correspond to (some of) the 9 actions of the experiment to maximize the variance between the 9 actions with regard to associated intentions and feelings. Questions 12-16 used a two-sided 6-point Likert scale and referred to more general mental state dimensions based on Tamir et al. (2016). Values reflect mean ratings ± standard deviations.

|  | rating question | open<br>pizza | open<br>bottle | open<br>present | close<br>pizza | close<br>bottle | close<br>present | squash<br>pizza | squash<br>bottle | squash<br>present |
| --- | --- | --- | --- | --- | --- | --- | --- | --- | --- | --- |
| 1 | satisfy a basic need? | 5.65 ± 0.72 | 5.81 ± 0.33 | 2.19 ± 1.18 | 3.35 ± 1.70 | 3.88 ± 1.75 | 1.73 ± 0.78 | 2.15 ± 1.16 | 2.08 ± 1.13 | 1.42 ± 0.76 |
| 2 | fulfil curiosity? | 2.62 ± 1.36 | 1.58 ± 0.61 | 5.23 ± 1.32 | 1.62 ± 1.02 | 1.38 ± 0.62 | 2.23 ± 1.48 | 1.50 ± 0.89 | 1.35 ± 0.43 | 1.38 ± 0.46 |
| 3 | make others happy? | 4.62 ± 1.34 | 3.77 ± 1.51 | 4.69 ± 0.95 | 3.15 ± 1.05 | 2.85 ± 1.16 | 4.31 ± 1.22 | 2.38 ± 1.14 | 2.19 ± 1.01 | 2.19 ± 1.01 |
| 4 | keep in order? | 1.50 ± 0.74 | 1.65 ± 0.69 | 2.08 ± 1.26 | 5.08 ± 0.93 | 4.73 ± 1.13 | 5.08 ± 0.93 | 5.15 ± 1.16 | 5.00 ± 1.15 | 3.88 ± 1.91 |
| 5 | store? | 1.58 ± 0.70 | 1.65 ± 0.66 | 2.08 ± 1.38 | 5.08 ± 1.34 | 4.23 ± 1.56 | 4.77 ± 1.48 | 3.15 ± 2.00 | 3.08 ± 1.90 | 2.38 ± 1.70 |
| 6 | do something self-rewarding? | 4.27 ± 1.13 | 3.62 ± 0.79 | 4.50 ± 1.12 | 2.54 ± 1.13 | 2.65 ± 1.09 | 3.88 ± 1.42 | 1.73 ± 0.63 | 1.88 ± 0.85 | 2.23 ± 1.03 |
| 7 | is s/he satisfied? | 4.96 ± 0.83 | 4.35 ± 0.88 | 4.58 ± 0.84 | 4.65 ± 1.07 | 4.08 ± 0.98 | 4.38 ± 1.12 | 4.08 ± 1.30 | 3.62 ± 1.37 | 3.23 ± 1.15 |
| 8 | is s/he excited? | 4.46 ± 1.09 | 2.85 ± 0.99 | 4.81 ± 0.88 | 2.62 ± 1.21 | 2.65 ± 0.97 | 4.23 ± 1.01 | 2.42 ± 1.13 | 2.62 ± 1.29 | 3.08 ± 1.15 |
| 9 | is s/he thankful? | 4.88 ± 1.16 | 3.85 ± 1.34 | 4.62 ± 1.10 | 4.04 ± 1.27 | 3.58 ± 1.12 | 3.96 ± 1.28 | 3.15 ± 1.16 | 2.73 ± 0.63 | 2.69 ± 1.15 |
| 10 | is s/he annoyed? | 1.46 ± 0.48 | 1.54 ± 0.63 | 1.54 ± 0.63 | 1.85 ± 0.80 | 2.08 ± 0.93 | 1.69 ± 0.90 | 2.31 ± 1.05 | 2.81 ± 1.22 | 3.15 ± 1.23 |
| 11 | is s/he regretful? | 1.54 ± 0.52 | 1.50 ± 0.46 | 1.50 ± 0.61 | 2.15 ± 0.88 | 1.65 ± 0.55 | 1.92 ± 0.93 | 2.04 ± 0.83 | 2.38 ± 1.31 | 2.92 ± 1.43 |
| 12 | positive or negative emotions? | 5.31 ± 0.95 | 4.27 ± 0.86 | 4.96 ± 0.85 | 4.15 ± 0.88 | 3.85 ± 0.83 | 4.50 ± 1.14 | 3.23 ± 0.78 | 3.04 ± 0.56 | 2.31 ± 1.01 |
| 13 | high or low arousal? | 4.77 ± 0.78 | 3.19 ± 0.85 | 4.69 ± 0.93 | 2.81 ± 0.75 | 2.62 ± 1.02 | 3.96 ± 0.95 | 2.65 ± 1.05 | 3.12 ± 1.14 | 3.35 ± 1.31 |
| 14 | rational or irrational? | 4.81 ± 1.13 | 4.73 ± 1.24 | 4.50 ± 1.34 | 4.62 ± 1.31 | 4.73 ± 1.15 | 4.96 ± 0.66 | 4.42 ± 1.46 | 4.12 ± 1.65 | 3.08 ± 1.50 |
| 15 | body or mind satisfaction? | 4.92 ± 1.13 | 5.31 ± 0.97 | 1.96 ± 1.14 | 4.31 ± 1.42 | 3.58 ± 1.22 | 1.81 ± 0.80 | 3.69 ± 1.28 | 2.73 ± 1.41 | 3.12 ± 1.31 |
| 16 | social or nonsocial? | 3.19 ± 1.65 | 2.38 ± 1.62 | 4.96 ± 1.30 | 2.77 ± 1.64 | 2.23 ± 1.44 | 4.54 ± 1.48 | 2.69 ± 1.65 | 2.50 ± 1.47 | 2.77 ± 1.64 |

**Supplementary Table 2:** Peak coordinates from main clusters of the crossmodal decoding in MNI space.

| Region | x | y | z | accuracy | t | p |
| --- | --- | --- | --- | --- | --- | --- |
| left MTG | -54 | -48 | -3 | 14.64 | 7.16 | 1.79E-07 |
| right MTG | 60 | -63 | 3 | 13.63 | 6.93 | 2.95E-07 |
| left aIPL | -63 | -30 | -33 | 14.54 | 8.06 | 2.60E-08 |
| right aIPL | 48 | -18 | -36 | 14.25 | 6.67 | 5.30E-07 |
| left PMC | -51 | 12 | 45 | 13.72 | 6.97 | 2.69E-07 |
| left VTC | -39 | -43 | -18 | 14.46 | 6.23 | 1.42E-06 |
| right VTC | 43 | -45 | -12 | 14.08 | 7.55 | 7.55E-08 |
| left PC | -3 | -60 | 42 | 14.65 | 8.38 | 1.36E-08 |
| left dmPFC | -12 | 39 | 33 | 13.78 | 7.00 | 2.53E-07 |
| left vLPFC | -57 | 36 | 3 | 13.98 | 7.49 | 8.59E-08 |
| left TPJ/pIPS | -33 | -81 | 42 | 14.38 | 8.52 | 1.01E-08 |
| right TPJ | 39 | -60 | 36 | 14.03 | 6.40 | 9.73E-07 |

